# SSMD: A semi-supervised approach for a robust cell type identification and deconvolution of mouse transcriptomics data

**DOI:** 10.1101/2020.09.22.309278

**Authors:** Xiaoyu Lu, Szu-Wei Tu, Wennan Chang, Changlin Wan, Jiashi Wang, Yong Zang, Baskar Ramdas, Reuben Kapur, Xiongbin Lu, Sha Cao, Chi Zhang

## Abstract

Deconvolution of mouse transcriptomic data is challenged by the fact that mouse models carry various genetic and physiological perturbations, making it questionable to assume fixed cell types and cell type marker genes for different dataset scenarios. We developed a **S**emi-**S**upervised **M**ouse data **D**econvolution (SSMD) method to study the mouse tissue microenvironment (TME). SSMD is featured by (i) a novel non-parametric method to discover data set specific cell type signature genes; (ii) a community detection approach for fixing cell types and their marker genes; (iii) a constrained matrix decomposition method to solve cell type relative proportions that is robust to diverse experimental platforms. In summary, SSMD addressed several key challenges in the deconvolution of mouse tissue data, including: (1) varied cell types and marker genes caused by highly divergent genotypic and phenotypic conditions of mouse experiment, (2) diverse experimental platforms of mouse transcriptomics data, (3) small sample size and limited training data source, and (4) capable to estimate the proportion of 35 cell types in blood, inflammatory, central nervous or hematopoietic systems. In silico and experimental validation of SSMD demonstrated its high sensitivity and accuracy in identifying (sub) cell types and predicting cell proportions comparing to state-of-the-arts methods. A user-friendly R package and a web server of SSMD are released via https://github.com/xiaoyulu95/SSMD.

**Key points:**

- We provide a novel tissue deconvolution method, namely SSMD, which is specifically designed for mouse data to handle the variations caused by different mouse strain, genetic and phenotypic background, and experimental platforms.
- SSMD is capable to detect data set and tissue microenvironment specific cell markers for more than 30 cell types in mouse blood, inflammatory tissue, cancer, and central nervous system.
- SSMD achieve much improved performance in estimating relative proportion of the cell types compared with state-of-the-art methods.
- The semi-supervised setting enables the application of SSMD on transcriptomics, DNA methylation and ATAC-seq data.
- A user friendly R package and a R shiny of SSMD based webserver are also developed.

## INTRODUCTION

The mouse has long served as the premier model organism for studying human biology and disease, due to their striking genetic homologies and physiological similarity to humans, as well as the relatively low cost of maintenance. Currently, thousands of unique inbred strains and genetically engineered mutants have been made available for a wide array of specific disease types [1]. Research on mouse models have provided added impetus and indispensable tool for studying human disease, regarding its initiation, maintenance, progression and response to treatment, as well as evaluating drug safety and efficacy [2] [3]. Amongst all, the ability to examine physiological states and interactions between diseased cells and their microenvironment in vivo represents the most important tool for studying disease dynamics. To this end, numerous omics data have been collected from mouse that vary in terms of genetic perturbations, cell/tissue types, and treatment conditions [4-7]. A strong computational capability is needed to study the interactions of components within the mouse tissue microenvironment subject to different genetic and physiological perturbations, the knowledge gained from which could be projected to human disease scenarios and provide invaluable insight and guidance for effective human therapeutic regimes.

Tissue transcriptomic data display convoluted signals from different cell types [8]. Deconvoluting cell components and identifying mouse strain-/tissue-/experimental condition-specific cell types and gene expressions are crucial for understanding how experimentally perturbed conditions are associated with cellular level characteristics and cell-cell interactions [9]. While multiple deconvolution methods have been developed for investigating the heterogeneous cell types in human cancer or other tissues data [10-19], they may not be directly applicable to mouse tissue data. First of all, the cell type specific genes for human cells differ from mouse cells; secondly, compared with human, the variations among different mouse tissue samples may be considerably higher, as they are collected from different strains with varied genetic background and experimental conditions.

Currently, ImmuCC and its varied versions are the only method specifically focusing on mouse data deconvolution [20]. The core computational algorithm, which was adapted from CIBERSORT designed for human [13], assumes fixed cell type and signatures gene expressions (subject to simple transformations) regardless of experimental conditions of the target data. This assumption becomes problematic as mouse data, which are collected from different strains, have varied genetic background, thus, it is expected the tissue compositions are highly adaptable regarding the existent cell types and their expression profiles [21-23]. Aside from prominent variability in the appearance of cell types and the expression levels of markers genes, mouse data deconvolution also suffers from the following challenges: diverse experimental platforms, prevalently small sample size of mouse experiments, and limited training data sets available for deriving signature genes of cell types.

To address these challenges, we developed a novel semi-supervised deconvolution method, namely Semi-Supervised Mouse data Deconvolution (SSMD), to infer data/tissue specific cell type marker genes and their expression profiles and estimate their relative abundances from transcriptomics data. SSMD is capable to infer the relative proportion of 35 cell types in the blood, inflammatory, cancer, central nervous system and hematopoietic system. To the best of our knowledge, SSMD is the only mouse data deconvolution method considering strain, tissue type and data specificity of cell type specific gene markers. We demonstrated SSMD achieved a high sensitivity in identifying the appearance of immune and stromal cell types in inflammatory tissue and brain cell types in central nervous tissue, and with a high accuracy in estimating their relative proportion on single cell RNA-seq simulated bulk tissue data sets. We also experimentally validated that the cell populations inferred by SSMD accurately recapitulates the true cell proportions measured by fluorescence-activated cell sorting (FACS) on a leukemia bone marrow data. Applications of SSMD on a large collection of public mouse blood, brain, cancer, and other inflammatory tissue data suggested that the method achieved a robust performance throughout diverse types of experimental conditions and platforms including RNA-seq, microarray and immuno-assay. In addition, the software of SSMD grants users to build in their own tissue/data specific knowledge of cell type specific markers to reinforce the method. An R package of SSMD is released through GitHub: https://github.com/xiaoyulu95/SSMD and a R Shiny based web server of SSMD is available at https://ssmd.ccbb.iupui.edu/.

## RESULTS

### Mathematical consideration and problem formulation

Denote 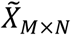 as a tissue data of *M* genes and *N* samples, a deconvolution analysis assumes 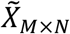 as the following non-negative product form:

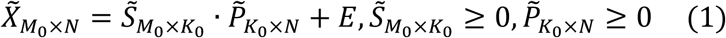

Here, 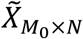 represents the observed gene expression matrix of *M*_0_ selected genes (a subset in *M*) in *N* tissue samples, and columns in 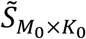, and rows in 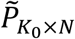, denote the expression signatures, and the relative proportions of the *K*_0_ cell types respectively. In the conventional formulation of deconvolution analysis, with fixed *M*_0_ and *K*_0_, 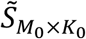 and 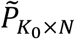 are solved to minimize the ℒ_2_ loss of the above linear equation. Because of the highly varied genetic and phenotypic background of mouse experiment, 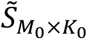, *M*_0_ and *K*_0_ are usually varied and unknown, i.e. for each 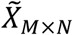 collected from tissues of certain microenvironment, what cell types are present, what gene markers each cell type expresses and how much they were expressed, could vary drastically due to the genetic and physiological perturbations. Correctly specified cell types *K*_0_, and selected cell type marker genes *M*_0_ can largely increase the prediction accuracy of 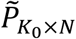. **Table 1** lists the key mathematical definitions utilized in this study.

**Table 1.**
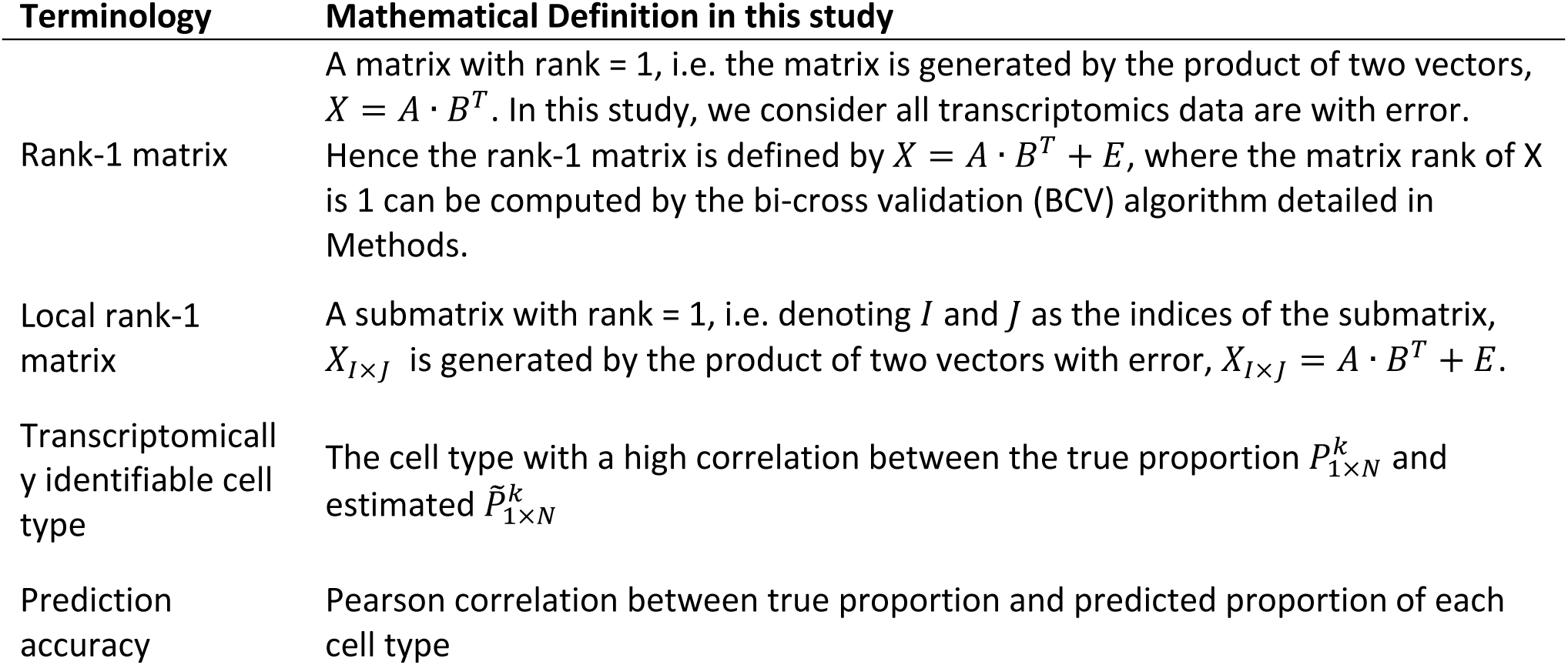
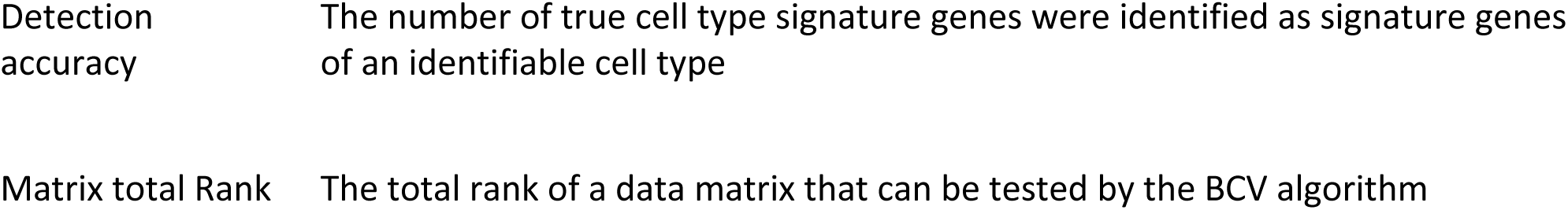
Definition of mathematical terms

In this study, we define a cell type *k* is “transcriptomically identifiable” if its ground-truth proportion 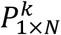 and estimated as 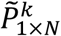 have high correlation, i.e.. 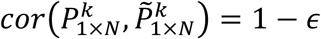 and *ϵ* is substantially small, where 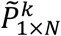 is the *k*th row of 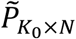, and *K*_0_ as the number of “identifiable” cell types. A strong condition for a cell type to be identifiable is that it has uniquely expressed genes [24]. Here we provided a comprehensive mathematical derivation of the relationship between cell type unique expression and identifiability of cell proportion in the **Supplementary Notes**. We derived the identity of cell type uniquely expressed gene markers, denoted as the set *G*_*k*_, is a necessary but non-sufficient condition for the identifiability of cell type *k*: – if *k* is “transcriptomically identifiable”, 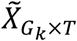 must be a matrix of rank one, for ∀ *T ⊂* {1, …, *N*}. This condition forms the foundation of how SSMD discover cell type marker genes that are not fixed, but instead specific to each dataset. Fortunately, we do not need to scan for all the local rank-1 matrices within 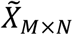, where *M* is usually to the tens of thousands. In fact, with an effective knowledge transfer of the gene labels derived from single or bulk cell training data, the genes that are more likely to be cell type specific markers of identifiable cell types can be detected, which forms the core algorithm of SSMD pipeline.

### SSMD Analysis pipeline

SSMD is a semi-supervised method composed by (1) training a large candidate list of cell type specific marker genes, (2) evaluating the identifiability of each cell type and confirming their marker genes for each to-be-deconvolved data, and (3) estimating the proportion of each cell type.

The training step is to look for genes that are more likely to serve as cell type marker genes through different tissue types and data sets, named as core marker lists. Specifically, we identified the genes that are commonly over expressed in one cell type comparing to the others in bulk cell data and commonly form rank-1 matrices in tissue data, by using a very extensive set of training data sets collected from different mouse strains and tissue types (see details in Methods). **Fig 1A** illustrates the procedure of SSMD to construct cell type core marker lists. On the bulk cell training data, we adopted a random-walk based approach to detect genes that are significantly expressed in higher quantities in one or a few cell types, than others (see details in Methods). As a result, a labeling matrix that annotates cell type specifically expressed genes will be constructed, which forms the first evidence of the potential marker genes for each cell type. Then on each bulk training tissue dataset, we further identified marker genes that form rank-1 submatrices with a community detection approach as detailed in methods. Only those modules, whose genes significantly and consistently over-represent one and only one cell type across multiple training tissue datasets, are selected to form the core marker list. Noted, variations caused by different experiment batches, tissue types and mouse strains were handled by enabling certain errors in the random-walk based cell type specific marker identification, i.e. identifying the genes overly expressed in the cell type comparing to the others in a certain proportion of the collected bulk cell data. In addition, data batch variation was also considered in the bulk data based training step, by identifying the genes commonly serve as cell type specific marker in more than 50% of analyzed bulk tissue training data. The goal of this training procedure is to summarize a relatively large list of commonly observed cell type specific marker genes, which can be used to as semi-supervised information to identify data set specific cell type marker for a further un-supervised deconvolution analysis.

**Fig 1.**
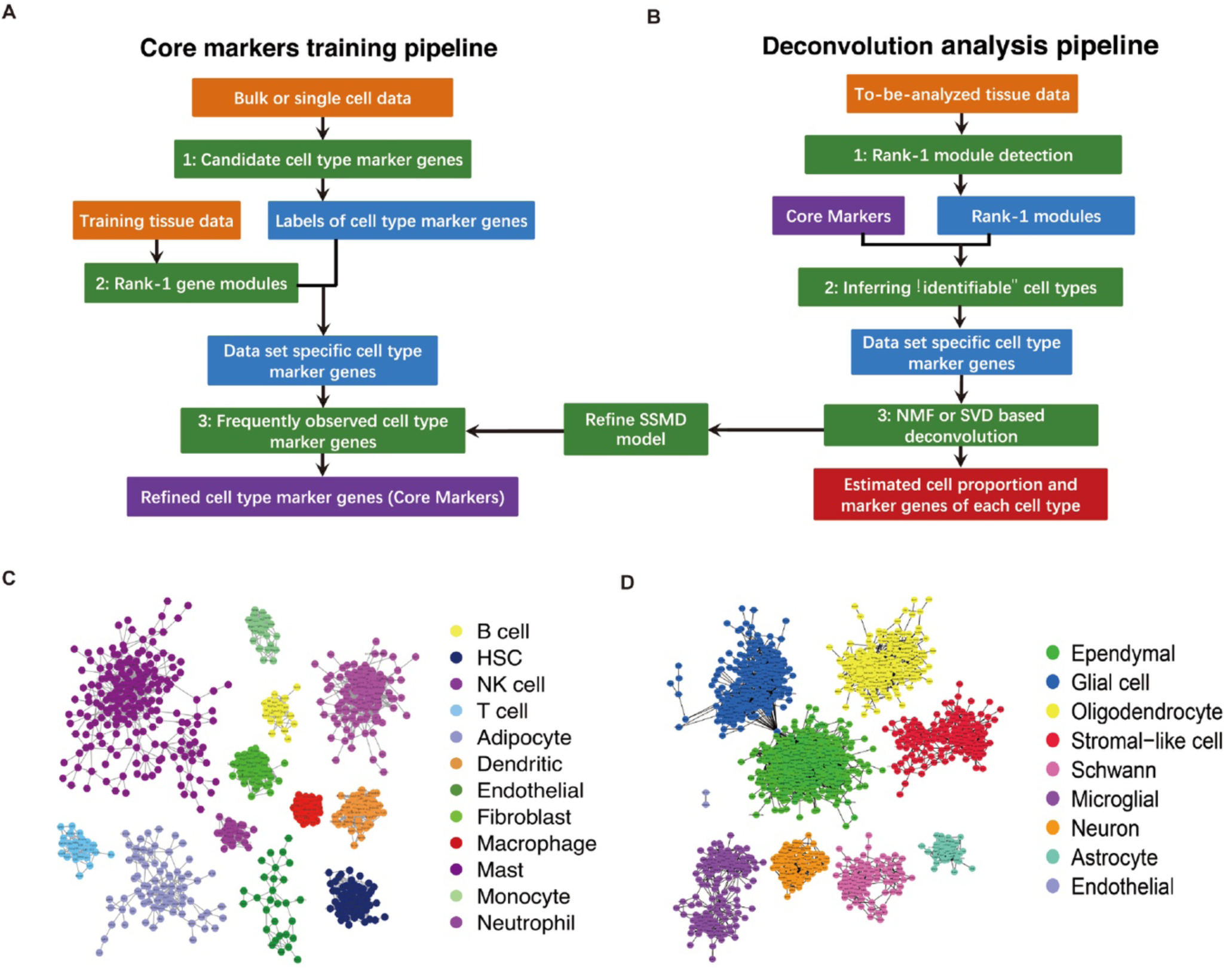
Analysis pipeline of SSMD and core cell type specific markers. (A) Analysis pipeline of the core marker training procedure. (B) Analysis pipeline of the deconvolution procedure. In (A) and (B), input data including training and target data, computational procedure and key intermediate outputs were colored by orange, green and blue, respectively. (C) Core markers of 12 cell types in blood, solid cancer, and inflammatory tissue. An edge between two genes means the two genes are co-identified as markers of one cell type in more than 50% of the training data sets. (D) Core markers of 9 cell types in central nervous system. Noted, core markers for the endothelial cell in the inflammatory tissue and central nervous system were separately trained by comparing with other cell types in the same tissue system.

Based on the cell type core markers, the deconvolution of any given bulk tissue dataset is composed by the steps as illustrated in **Fig 1B**. SSMD first identifies all the rank-1 modules on the target dataset by an iterative hierarchical clustering and bi-cross validation approach. Then SSMD selects the rank-1 modules that are likely to be markers of a certain cell type for this data set, if genes in the modules largely overlap with the core marker list of one and only one cell type. Modules that are highly co-linear will be merged. Consequently, genes in each module is called gene markers of one cell type, that satisfy the necessary condition for “transcriptomically identifiable”. Notably, two modules may represent the same cell type, and they are treated as marker genes of different subtypes of the cell type. Here, the total number of modules is an estimate of the number of “identifiable” cell types, i.e., *K*_0_. Importantly, SSMD is an “semi-supervised” approach, because the cell marker genes do not solely depend on the training data, but also the co-expression patterns of the marker genes in the target dataset. In other words, SSMD addresses the variability issue of signature genes from one dataset to another, and has the potential to discover cell types not pre-defined. Algorithms of each computational step are detailed in Materials and Methods. Complete flow chart of the SSMD pipeline is provided in **Supplementary Fig S1**.

The prediction of the cell type proportions is conducted using a constrained Non-negative Matrix Factorization (NMF) method by solving the following optimization problem:

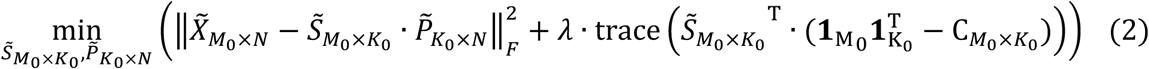

where 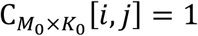 if gene *i* is marker of the cell type *j*, and 0 otherwise. **1**_d_ denotes an all-1 column vector of length *d, λ* is a hyperparameter selected by cross validation, and other annotations follow equation (1). The constraint matrix 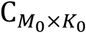 is enforced upon the regular NMF to guarantee similarity of the solved signature matrix 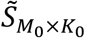 and constraint 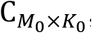, namely, in the *k*th column of 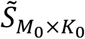, it should have higher expressions for genes that are markers of cell type *k*. The solution to (2) is by alternative update where each time one of 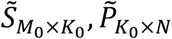 is held fixed, and the other is updated. *λ* can be tuned by using simulated tissue data with known cell proportion. In this study, we tuned *λ* and empirically select *λ* as 10 when 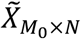 is log normalized microarray data or log(X+1) normalized FPKM/CPM/TPM RNA-seq data.

Following these procedures, and on a large collection of mouse bulk cell and tissue training data, we generated core marker gene lists for different tissue microenvironments: (1) for mouse blood, solid cancer and inflammatory tissues, 980 genes of 12 cell types namely T cell, B cell, NK cell, hematopoietic stem cell, monocyte, macrophage, neutrophil, mast cell, adipocytes, fibroblast, dendritic cell, and endothelial cell were discovered (**Fig 1C**); (2) for mouse hematopoietic system, 2877 genes of 14 cell types namely hematopoietic stem cell, common lymphoid progenitor, granulocyte-macrophage progenitors, megakaryocyte lineage-committed progenitor, erythroid cell, megakaryocyte-erythrocyte progenitors, multipotent progenitors, early myeloid progenitor, mature myeloid cell, pre colony forming unit erythroid, pre-megakaryocytic/erythroid progenitor, B cell, CD4+ T and CD8+ T cell were discovered (**Supplementary Table S1**), and (3) for mouse central nervous system tissue, 1570 genes of nine cell types namely ependymal cell, general glial cell, oligodendrocyte, stromal-like cell, Schwann cell, microglial, neuron, and astrocyte were discovered (Fig 1D). Complete lists of the core marker genes are given in **Supplementary Table S1**. It is noteworthy that the size of core marker list ranges from 27 to 547 for different cell types. However, our analysis suggested that more than 5 marker genes that form a rank-1 matrix is sufficient for an accurate estimation of cell proportion. Note that, compared with conventional regression based deconvolution analysis, SSMD only uses labels of the core markers as the semi-supervised information and identifies data set specific cell type markers for a further unsupervised estimation of cell types, which grants a flexibility and robustness to handle the variation of cell type specific marker genes and their expression scale through different mouse strains, tissue types and experimental platforms. In addition, the semi-supervised formulation of SSMD enables the inference of identifiability of each cell type and identification of rare or sub cell types.

### Benchmarking based on artificial tissue data simulated by using single cell RNAseq data

We first benchmarked SSMD on a set of artificial tissue data simulated from four single cell RNAseq (scRNA-seq) datasets of mouse lung, pancreas, small intestine and melanoma. For each data set, we simulated 100 tissue samples by randomly drawing and mixing cells of different types whose proportions follow random Dirichlet distributions. Prediction accuracy of each cell type was assessed by the Pearson correlation coefficients between its known mixing cell proportions and the predicted relative proportion. We compared SSMD with three state-of-arts deconvolution methods of mouse data, namely ImmuCC (ICC), tissue-ImmuCC (TICC) and EPIC [11]. Our analysis suggested that SSMD achieved 93.2% prediction accuracy on average in the four simulated data sets and 23 out of the 28 cell types (82.1%) are with higher than 0.9 prediction accuracy (**Fig 2A-D**). In contrast, EPIC, ICC and TICC achieved 69.7%, 45.2% and 48.5% averaged prediction accuracy on the cell types covered by these methods, and the proportion of cell types with higher than 0.9 prediction accuracy are 32.2% (9/28), 0% (0/28) and 7.2% (1/14), respectively. We also tested the popular human data deconvolution methods such as CIBERSORT (CIBERSORTx) and TIMER [9, 13], by using the known human and mouse homolog genes. Non-surprisingly, predictions made by CIBERSORT and TIMER on the mouse are less accurate than SSMD. TIMER and CIBERSORT achieved 49.25% and 47.5% averaged prediction accuracy, and the proportion of cell types with higher than 0.9 prediction accuracy are 17.9% (5/28) and 3.6% (1/28) (**Supplementary Table S4**).

**Fig 2.**
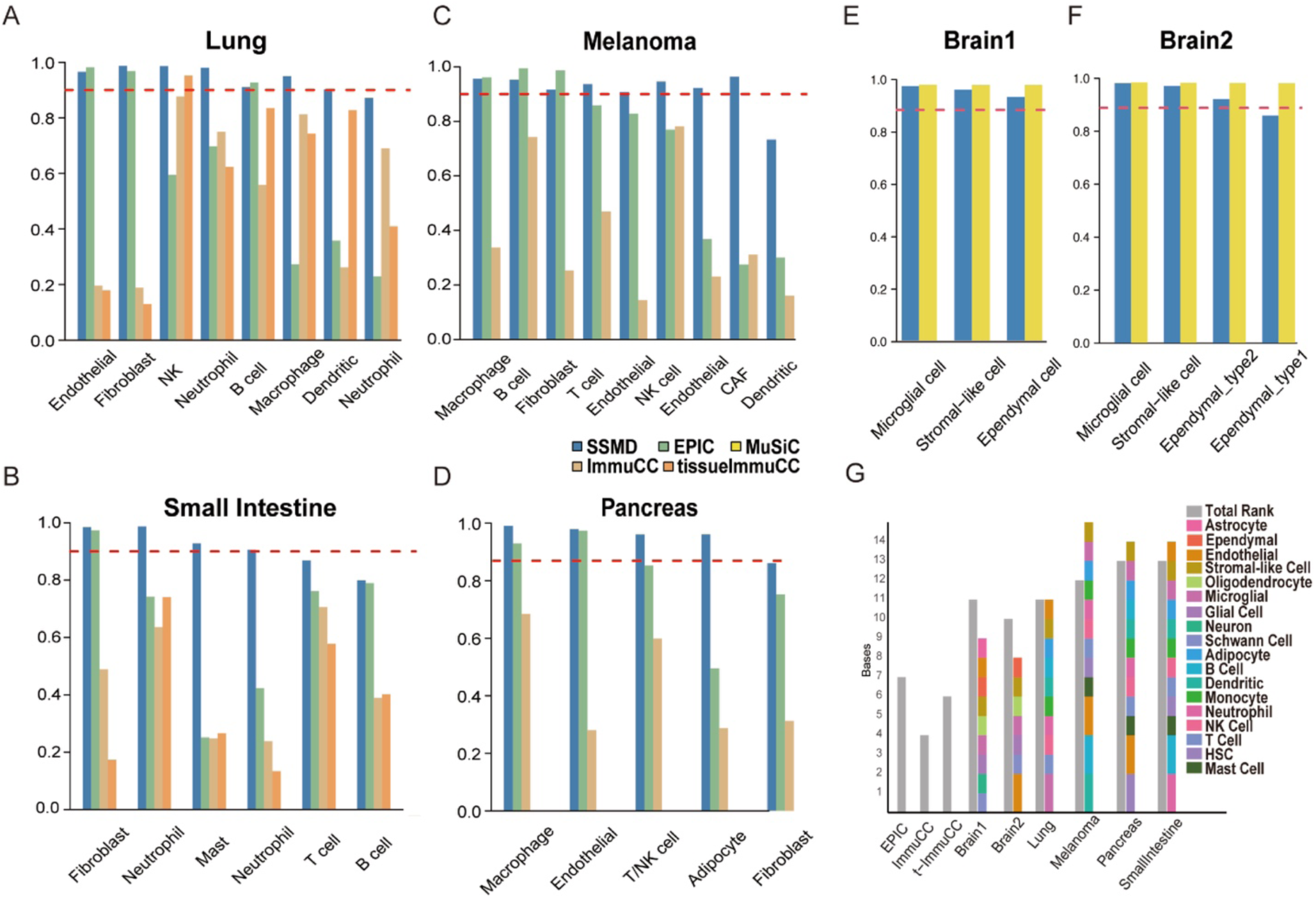
Method evaluation on scRNA-seq simulated tissue data. (A-D) Correlation between true and predicted cell proportions in the simulated Lung (A), Pancreas (B), Small Intestine (C), and Mouse Melanoma (D) tissue data. The x-axis represents cell type and y axis represents prediction accuracy. Predictions made by SSMD, EPIC, ImmuCC and tissue-ImmuCC were dark blue, green, yellow and orange colored, respectively. The red dash line represents the 0.9 correlation cutoff. (E-F) Correlation between true and predicted cell proportions in the two simulated brain tissue data. (G) The total rank of the gene expression profile of selected marker genes in the six simulated tissue data (grey), and the total number of cell types identified by SSMD in each data set or assumed in other methods (left three grey bars).

It is noteworthy that the SSMD enables the detection of sub cell types defined as transcriptomically identifiable. SSMD successfully identified two sub populations of fibroblast cells in the melanoma data and different subtypes of neutrophils in lung and small intestine data. In contrast, ICC, TICC and EPIC are not capable of providing cell subtype predictions due to their fixed cell type assumption.

We also benchmarked SSMD on simulated brain tissue data using two scRNA-seq data of central nervous systems. SSMD achieved more than 0.9 correlation in predicting the cell types microglial, stromal-like, and ependymal subtypes in the simulated tissue data (**Fig 2E-F**). To the best of our knowledge, SSMD is the first of its kind method to specifically target mouse central nervous system decomposition. To benchmark SSMD, we selected MUSIC as the state-of-the-art method, which requires an additional input of an scRNA-seq data to train context specific gene signatures [25]. Here we first utilized the same scRNA-seq data for tissue data simulation and signature training in MUSIC. Non-surprisingly, MUSIC achieved consistently good predictions (averaged cor=0.99), and the predictions made by SSMD are very close to MUSIC with slightly lower correlations compared with MUSIC under this ideal setup. In sight the possible disparity caused by tissue, strain, and experimental platform variations between the target tissue data and available scRNA-seq data for training cell markers, we also conducted a robustness test of MUSIC and SSMD (see details in Supplementary Notes). Our analysis suggested that MUSIC highly depends on the consistency of cell type specific marker genes and their expression scale between the target tissue and the training scRNA-seq data. In contrast, the de novo data set specific marker identification by SSMD enables a broader application to the tissue data without matched scRNA-seq data. Because EPIC, ImmuCC and tissue-ImmuCC cannot analyze brain tissue data and the melanoma and pancreas tissue were not covered by tissue-ImmuCC, we did not include the comparison with these methods on the brain tissue data.

To further validate the specificity of SSMD, we tested the total rank of the identified marker genes and compared with the number identified cell types (TIMER and CIBERSORT achieved 49.25% and 47.5% averaged prediction accuracy. and the proportion of cell types with higher than 0.9 prediction accuracy are 17.9% (5/28), and 3.6% (1/28).). We also compare the total matrix rank of the marker genes used in other methods and the number of cell types assumed in those methods. Comparing to the fixed number of cell types in other methods, the number of cell types predicted by SSMD better matches the total rank of the expression profile of identified marker genes. Our observation suggested SSMD can correctly estimate the number of cell types and select proper markers for cell type proportion estimation. It is noteworthy the predicted number of cell types may not exactly match the total rank of selected markers because possible co-linearity among the true proportion of the cell types.

### Experimental validation of SSMD by using matched RNA-seq and cell sorting data

We generated a tissue RNA-seq data of 11 mouse bone marrow tissue samples with matched cell counting using Fluorescence activated cell sorting (FACS) (see details in Methods). Application of SSMD on the RNA-seq data identified hematopoietic stem cell (HSC), general myeloid progenitor (GMP), mature myeloid cell and Pre-B cells, and their cell type specific markers. We also observed that the correlation between SSMD predicted and FACS measured amount of HSC, GMP, mature myeloid cell and B cells are 0.92, 0.8, 0.86, and 0.97, respectively, suggesting a high prediction accuracy of SSMD. **Fig 3A-D** shows the correlation between the SSMD predicted cell proportion and the FACS measured cell proportion of the four cell types. **Fig 3E-H** illustrate the FACS based cell counting of the four cell types. Complete cell type specific markers, cell proportions counted by FACS and predicted by SSMD were given in **Supplementary Table S2**. It is noteworthy that SSMD is not compared with other methods as none of the existing method is capable of predicting proportions of hematopoietic cell types.

**Fig 3.**
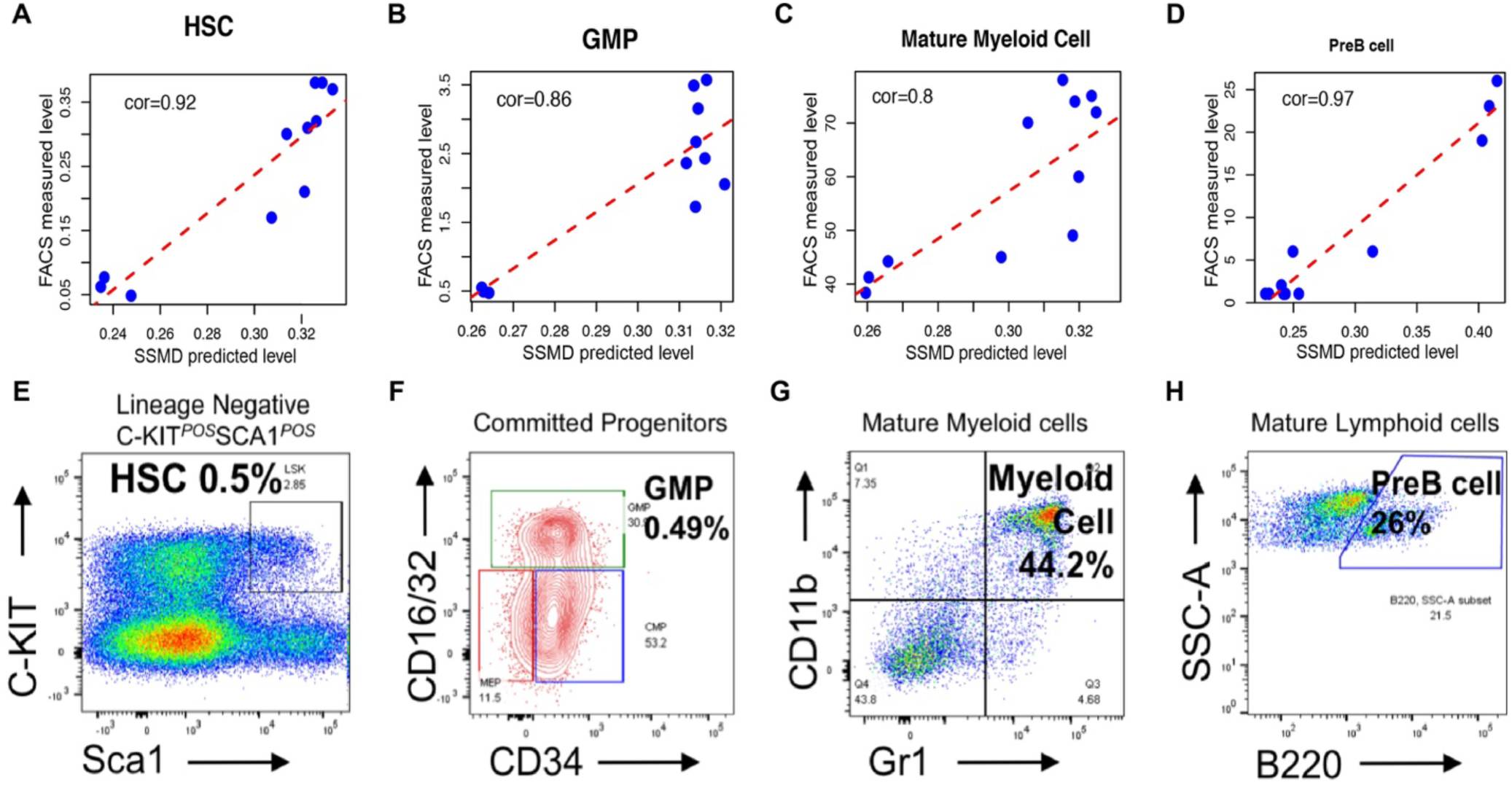
Method evaluation on scRNA-seq simulated tissue data on hematopoietic tissue data. (A-D) Correlation between SSMD predicted (x-axis) and FACS identified (y-axis) cell proportions of HSC, GMP, mature myeloid cell and preB cell. (E-H) marker proteins utilized to identify the four cell types by using FACS. The x- and y-axis of the plots represent the level of cell type markers. The black block in (E), the green block in (F), the upper-right block in (G) and the block in (H) are the sorted HSC, GMP, Myeloid and Pre-B cell, respectively.

### Application of SSMD to real mouse tissue transcriptomics data

We applied SSMD to nine cancer and eight central nervous system tissue data of four different experimental platforms, including one data set measured by immune-assay. On average, SSMD identified more than seven cell types in each of the cancer data, and the number of identified cell types is highly consistent with the total rank of the expression profile of the detected cell type specific marker genes (**Fig 4A**). This indicates that SSMD is capable of capturing the latent structure of the data. We further examined the explanation score (E-score), defined as the averaged absolute residual of the non-negative linear regression of each marker gene’s expression on the predicted cell proportion, i.e. the average measure of how the predicted proportions could explain all the marker genes’ expression levels. A high E-score is a necessary condition for an accurate cell proportion prediction. On average, the data set specific markers genes of each cell type identified by SSMD achieved 0.73 E-score while the average E-score of the marker genes used by EPIC and ImmuCC is 0.45 and 0.3 (**Fig 4B**). Similarly, application of SSMD on eight central nervous system tissue data identified more than seven cell types on average. The number of identified cell types is highly consistent with the total rank of the gene expression profile of the marker genes (**Fig 4C**). And the marker genes identified by SSMD achieved averaged 0.77 E-score for the cell types in central nervous system (**Fig 4D**). It is noteworthy that multiple marker sets of fibroblasts, myeloid or microglial cells that forming distinct rank-1 bases were identified in numerous data sets, suggesting the possible sub types of these cell types identified by SSMD.

**Fig 4.**
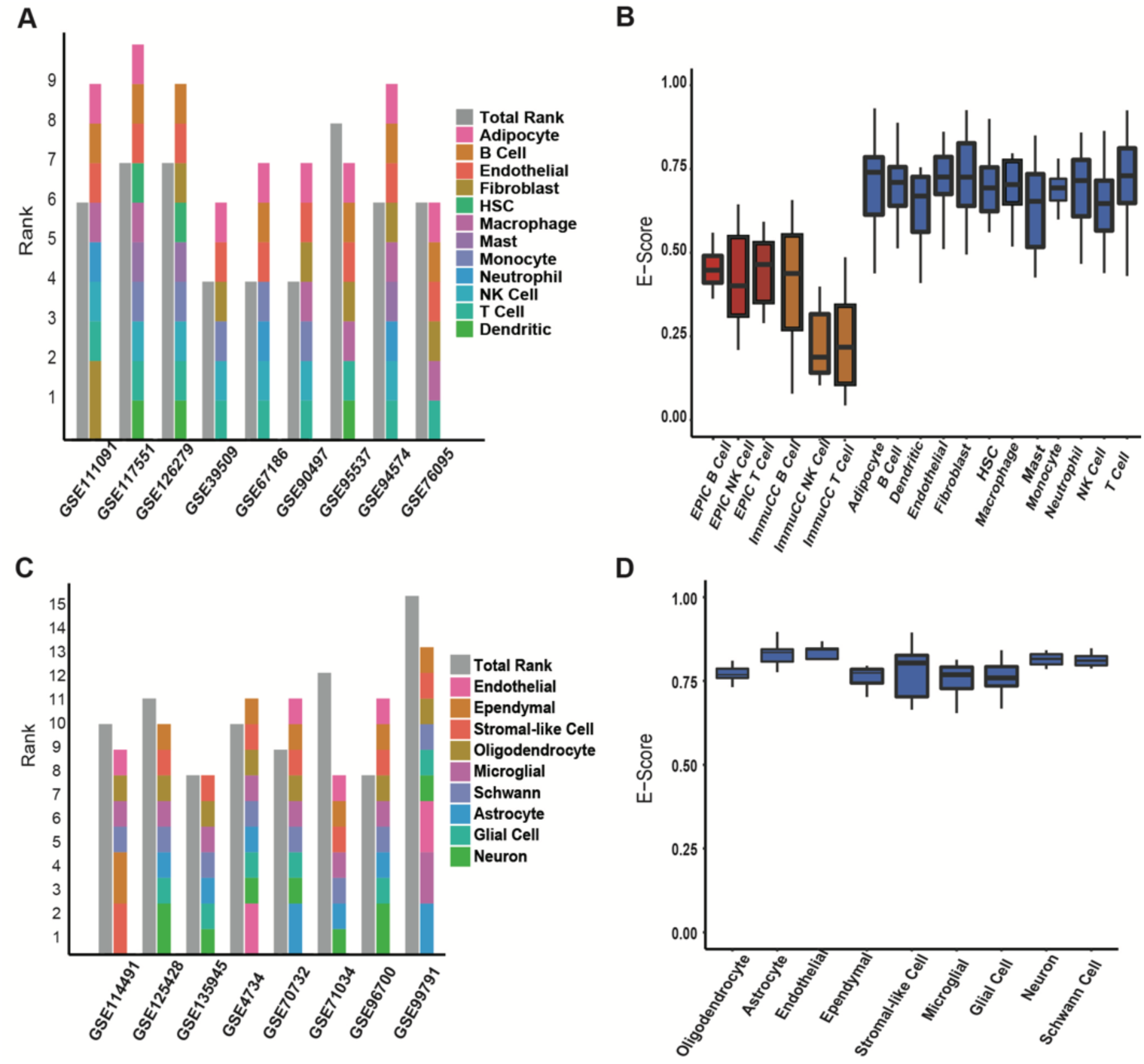
Prediction of SSMD on real tissue data. (A, C) The total rank of the gene expression profile of selected marker genes (grey) in different (A) cancer tissue and (C) brain data, and the total number of cell types identified by SSMD in each data set (colored). (B, D) E-Score for different cell types identified by SSMD (blue) in (B) cancer and (D) brain data set or assumed in other methods (EPIC: red, ImmuCC: Yellow).

### Robustness analysis

We first evaluated the variation of cell type specific markers through different mouse strains on one transcriptomic dataset of mouse liver tissue samples collected from 31 different mouse strains [26]. To the best of our knowledge, this is the only dataset in the public domain that systematically measured gene expression profiles of the same tissue type for different mouse strains by using the same experimental platform. SSMD was applied to the data of each mouse strain respectively. 9 cell and their sub types were commonly identified in the liver tissue of most strains. The identifiability of the cell types and the detected cell type markers among different strains were compared (**Fig 5**). We analyzed all the identified marker genes that form rank-1 modules, i.e. the necessary condition for gene markers of identifiable cell types, and noticed that only 9.1% of the identified marker genes are shared in more than 50% strains, while 58.4% of the identified marker genes only served as a cell type marker in less than 20% of the analyzed strains, suggesting a high variation of cell type specific markers among different mouse strains, and the necessity to consider strain or data set specificity in deconvolution analysis.

**Fig 5.**
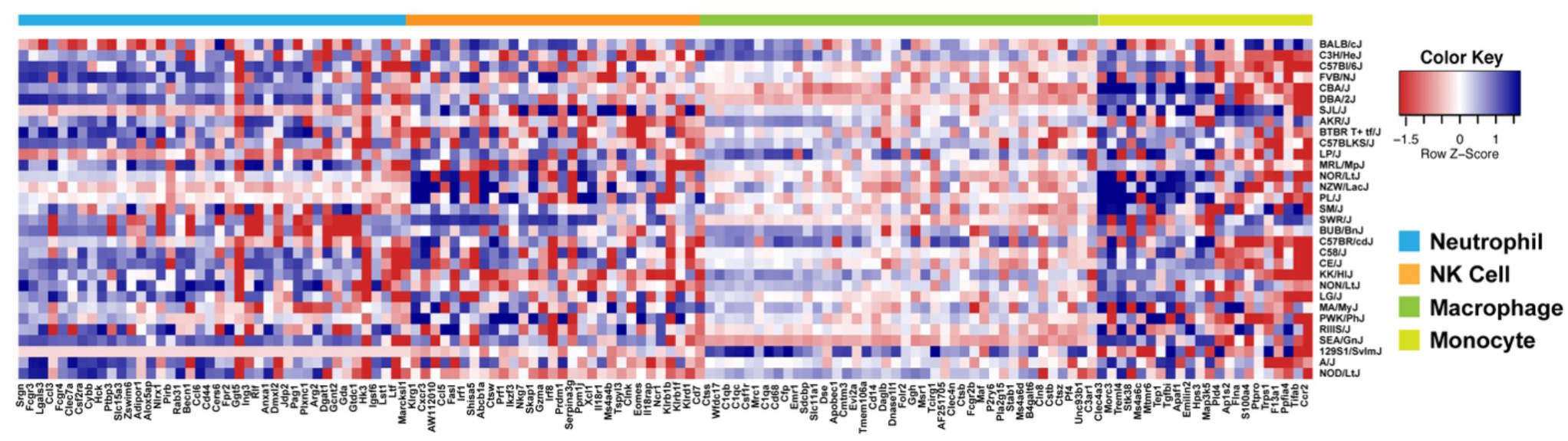
Correlation between expression level of strain specific cell type marker genes and predicted cell proportion. High correlation is a necessary but non-sufficient condition for the genes to serve as marker genes of the cell types in corresponding mouse strain. In the heatmap, *x*- and *y*-axis represent genes and mouse strains, respectively. Genes in the core marker list of four selected cell types, namely Neutrophil, Nature Kill (NK), Macrophage, and Monocyte, were colored on the column side bar.

We further examined the robustness of SSMD by evaluating its (1) sensitivity and (2) specificity in identifying cell types specific marker genes and its (3) accuracy in assessing of cell proportions, on the data of different sample sizes. Previous studies revealed that the robustness of the computation of co-expression correlation will decrease when the sample size is below 25. To comprehensively evaluate the method’s robustness, we selected five data sets, namely GSE76095, GSE67186, GSE90885, GSE94574, and GSE126279, with sample size ranging from 15 to 30 and randomly drew samples from each data set to build testing data sets of different sample size. We assumed the cell type markers and cell proportion inferred from whole data as “true” markers and proportions, and evaluated the consistency between the “true” ones and the ones predicted from small sub data sets. Accuracy in cell proportion prediction was assessed by the Pearson correlation between proportions predicted from small data and the “true” proportion on overlapped samples.

On average, all of the marker genes of the “true” cell types were also identified when sample size is low (**Fig 6A**). In addition, the cell proportion of 92.3%, 94.6% and 98.9% of the correctly identified cell types were with more than 0.9 correlation with their “true” proportions when the sample size is 6, 12 and above 20 (**Fig 6A**). Our analysis suggested a high robustness of the sensitivity and prediction accuracy of SSMD when sample size is as small as 6, i.e. the commonly used sample size in two-condition-comparison experiment (3 samples vs 3 samples). However, as a trade-off, there is a high false discovery rate of cell type specific modules when sample size is small, due to the low specificity of gene co-express analysis. To control the false discoveries on small data sets, we further derived a more “stringent” set of 341 cell type specific marker genes among the core marker set (see details in Methods). Our method validation demonstrated a slight drop of the sensitivity and prediction accuracy when using the stringent marker set on small data set (**Fig 6B**), while the specificity of the identified cell type specific markers increased to from 54.4% to 72.6% when sample size is above 12 (**Fig 6C**). **Fig 6D** illustrates the E-score of the cell type specific marker genes identified by using the core and the more stringent marker set with respect to different sample size. The E-score of the cell types marker genes identified by using the more stringent marker set were significantly higher than the ones identified by using the general core marker sets when sample size is below 10, also demonstrating the stringent core marker sets can effectively increase the analysis specificity when sample size is small.

**Fig 6.**
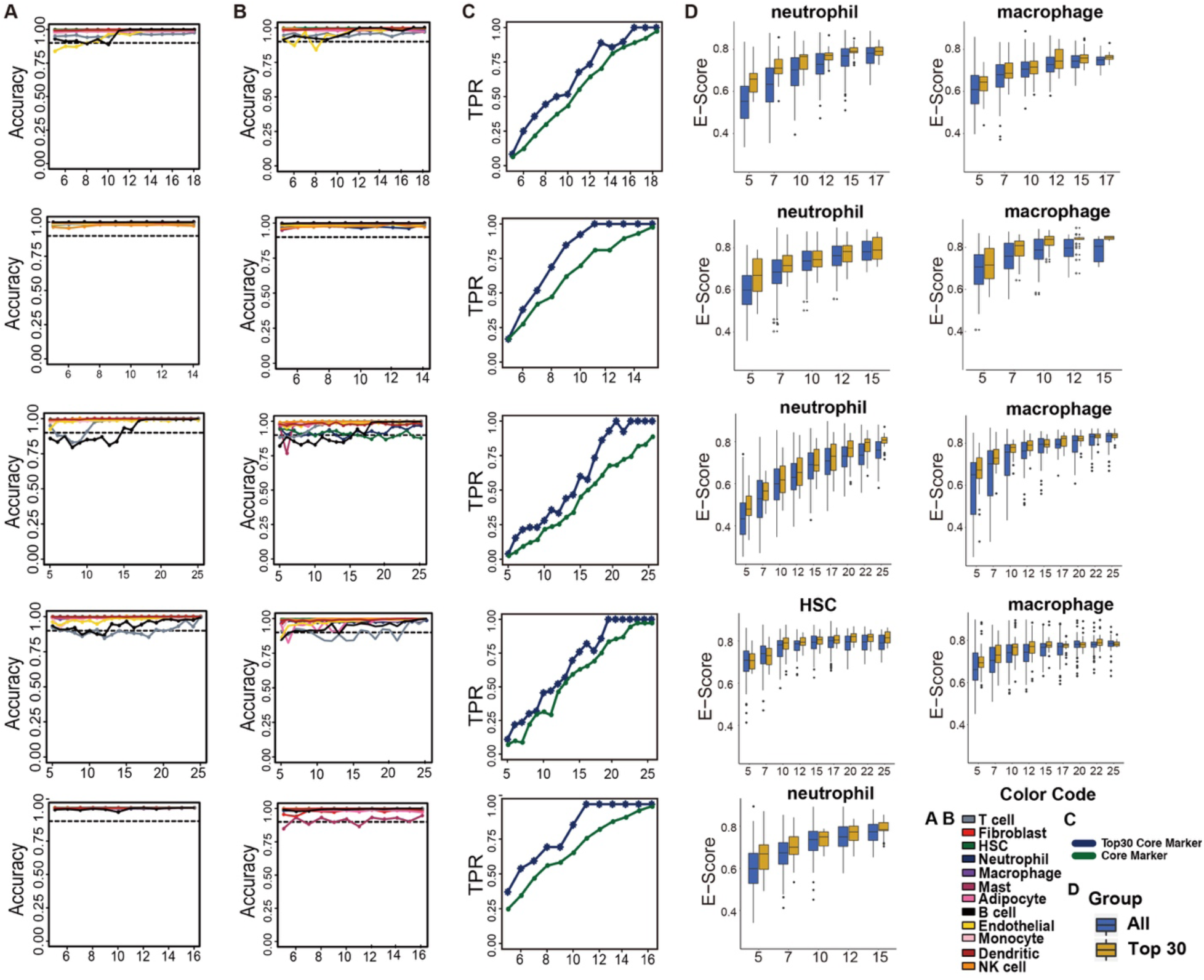
Performance evaluation of different sample size. (A) Prediction accuracy (*y*-axis) in different sample size (x-axis) using all core markers. Accuracy is the Pearson correlation between predicted proportion using only selected small sample and using all samples. (B) Prediction accuracy (y-axis) in different sample size (x-axis) using selected stringent markers. (C) True positive rate (*y*-axis) of the cell type specific markers identified by using the stringent markers (blue) and core markers (green) with respect to different sample size (x-axis). (D) E-Score for using co-expression modules consisting of all core markers and only selected stringent markers. From top to bottom, the statistics were derived from GSE76095, GSE67186, GSE90885, GSE94574, and GSE126279.

## DISCUSSION

Over the years, research using well-established mouse models to mimic human conditions have provided extensive insight into the mechanisms underlying many human diseases. We developed SSMD to study mouse tissue microenvironment of complex traits, to mine the interactions of cell components in the microenvironment, which will feed back to studying human microenvironment. In order to have a robust prediction of cell component abundance in mouse tissue, SSMD detects a subset of the genes and identifiable cell types that are the most representative to the tissues to be analyzed, instead of using fixed gene signatures and cell types as in classic deconvolution schemes. The limitation in expression profiling and the intrinsic and mysterious variability in microenvironments excludes the possibility to have a unified set of cell type specific genes that have absolutely constant expression across all conditions. The way SSMD flexibly defines cell type marker genes mitigates the impact of variable marker genes due to experimental platforms and microenvironment alterations. This strategy allows our model to fully recapitulate the disparity of cell types and their marker genes across different microenvironment and data-generating platforms. In addition, the semi-supervised formulation enables the detection of sub cell types, which has been validated on scRNA-seq data simulated tissue data. Hence, a relatively coarse standard for categorizing the cell types was used in training the core marker list, which enabled a high robustness of the core markers. The unsupervised constrained-NMF or SVD-based deconvolution on the selected marker genes further excludes the adversarial batch effects.

It is noteworthy a successful identification of the rank-1 modules depends on a relatively large samples (>25) sharing cell types and marker genes. Currently, SSMD cannot be applied to the data with a single or small sample size. However, we consider such a tradeoff between sample size and prediction robustness is highly worthwhile, especially considering using SSMD as an exploratory tool in large scale publicly available mouse transcriptomics data. After all, the predicted proportions are often to be associated with other biological and clinical features, which will be severely underpowered with a small sample size.

We released a R package of SSMD via https://github.com/xiaoyulu95/SSMD and a web server via https://ssmd.ccbb.iupui.edu/. As illustrated in **Supplementary Fig S2A**, the input data is a mouse tissue transcriptomics data and user selected tissue specific cell type core marker sets. Currently, SSMD offers general core and stringent marker sets of 6 cell types in blood system, 12 cell types in normal, inflammatory and cancer tissue, 9 cell types in central nerve systems, and 14 cell types in hematopoietic systems. **Supplementary Fig S2B** illustrates a practical guide for using SSMD of different tissues and sample size. The input of SSMD is a mouse tissue expression data set and user selected tissue environment category. The output of SSMD includes the identified data set specific cell type markers and the estimated sample-wise relative proportion of each identifiable cell type. We consider the currently included cell types are comprehensive enough to cover major cell types in mouse. However, the tissue specific cell types (for example, liver cells in liver tissue, colon cells in colon tissue, etc) were not included in our training scope. As forming rank-1 pattern among marker genes is a necessary but non-sufficient condition of identifiable cell types, SSMD R package can also output rank-1 modules that do not enrich the core markers of any cell type, which could possibly be markers of rare cell types. The user could further investigate whether the gene module corresponds to a real cell type or not. Another key feature of the webserver is that users are welcome to contribute their data to reinforce the training of cell type specific marker genes.

Potential future directions of SSMD include (1) enabling identification of cell type specific varied functions, which is not generally available for tissue data analysis in the public domain, (2) identifying data set specific cell type markers forming rank-1 submatrix in a subset of samples, i.e. local rank-1 submatrix, which can benefit from state-of-the-arts subspace clustering methods [27-29] and (3) extending and implementing the semi-supervised framework of SSMD with other state-of-the-arts deconvolution methods by refining data set specific cell marker genes. We anticipate that our computational concept, which is to identify data set specific and computationally “identifiable” cell types and their marker genes, can provide high robustness in deconvolution analysis, by which the predicted cell proportions can be reliably correlated with experimental features to provide biologically meaningful interpretation of the roles of microenvironmental changes in different disease tissues.

## MATERIALS AND METHODS

### Random walk based identification of cell type specifically expressed genes from tissue data

We applied a non-parametric random walk based approach to screen genes with higher expression in certain cell types comparing to others, using bulk cell training data. On the combined expression matrix containing M genes for N samples of K cell types, we first calculated the expected frequency of each cell type, i.e. dividing the total number of samples for the cell type (*N*_*k*_, *k* = 1, …, *K*) by the total number of samples N, denoted as *E*_*k*_ = *N*_*k*_/*N, k* = 1, …, *K*. For a given gene *g*, denote *x* and *x*^*k*^ as vectors of expression profile for cells of all types and type *k*. Denote *0*_*jk*_ as the percentage of values in *x*^*k*^ that are no less than the *j*th largest value in vector *x*. A random walk vector d_1×*N*_ that describes the non-negative discrepancy between the observed and expected cell type frequency of the gene was defined as 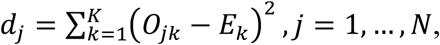 which attains a minimum value of zero at N. A higher peak of the random walk d_1×*N*_ suggests gene *g* is more enriched in certain cell types than the others. Denote *m* as the index of the maximum of d1×N, i.e. m = argmax (d_1×*N*_), and the cell type frequency at *m* as 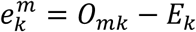 Cell types were further ordered by 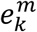 decreasingly, and a labeling matrix *L* was built such that 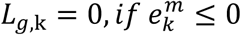 otherwise, 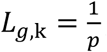, *if **x***^*k*^ has the pth largest mean among *x*^1,^ …, *x*^*k*^.

It is noteworthy the approach can be directly applied to scRNA-seq data for marker training. In this study, due to the relatively limited availability of existing scRNA-seq data, especially the mouse strain and tissue type coverage, we generate core marker list purely by using bulk cell data.

### Identification of rank-1 cell type uniquely expressed gene modules

To screen genes that form tight rank-1 modules on various tissue training datasets, SSMD performs a community detection method among the genes specifically expressed in each cell type as stored the labeling matrix. A correlation matrix was first built among cell type specifically expressed genes, and the significance cutoff of correlation was determined by random matrix theory. Random matrix theory (RMT) has been widely used to understand the low rank structure encoded in biological data. In this study, an RMT based approached developed by Luo et al was used to determine the threshold of significant correlation for each dataset[30]. rm.get.threshold functions in the RMThreshold R package was utilized. Specifically, RMT indicated that the nearest neighbor spacing distribution of eigenvalues will have a characteristic change when the threshold properly separates signal from noise. By removing all the below-threshold correlation elements, the co-expression modules can be more robustly unraveled. Then, hierarchical clustering was performed using the correlation matrix as similarity measure.

Specifically, SSMD gradually increases the height of the hierarchical clustering at which the tree is cut. At each height, the number of genes, the average correlation among the genes, and the rank of the matrix composed of the genes in each of the cluster, is calculated. Here, matrix rank is determined by a modified bi-cross validation (BCV) algorithm. SSMD stops scanning the hierarchical tree if all the clusters contain less than q_0_ genes, or the three following criterior is met for all the clusters: (1) with at least q_0_ genes, (2) the average correlation among the genes is above the threshold determined by RMT, and (3) the rank of the expression matrix profile of the genes in the cluster is 1. In this study, q_0_=7 is used. Such an iterative approach will eventually select the clusters with at least q_0_ genes, each of which is considered as possible cell specific marker genes specific to this data set. SSMD merges modules until the canonical correlation between any pair of module is lower than a cutoff *cor*_*cut*_ or the number of current modules is not larger than the total rank of the gene expression profile of the selected data set specific markers genes. In this study, we utilized *cor*_*cut*_ = 0.9.

### A modified Bi-cross validation rank test

Bi-cross validation (BCV) has been developed to estimate the matrix rank for singular value decomposition (SVD) and Non-negative Matrix Factorization (NMF), which requires a prefixed low dimension *K* and two low rank matrices for the approximation *X* _*M×N*_ = *W* _*M×K*_ · *H* _*K×N*_ The error distribution of gene expression data is usually non-identical/independent, mostly because a gene’s expression can be affected by its major transcriptional regulators, other biological pathways and experimental bias. Hence undesired biological characteristics and experimental bias may form significant dimensions in a gene expression data [31]. In sight of this, we developed a modified BCV rank test (*<u>Algorithm 1</u>*) to minimize the effect of the non-i.i.d errors in assessing the matrix rank of a gene expression data.

#### Algorithm 1: Modified Bi-cross validation matrix rank test

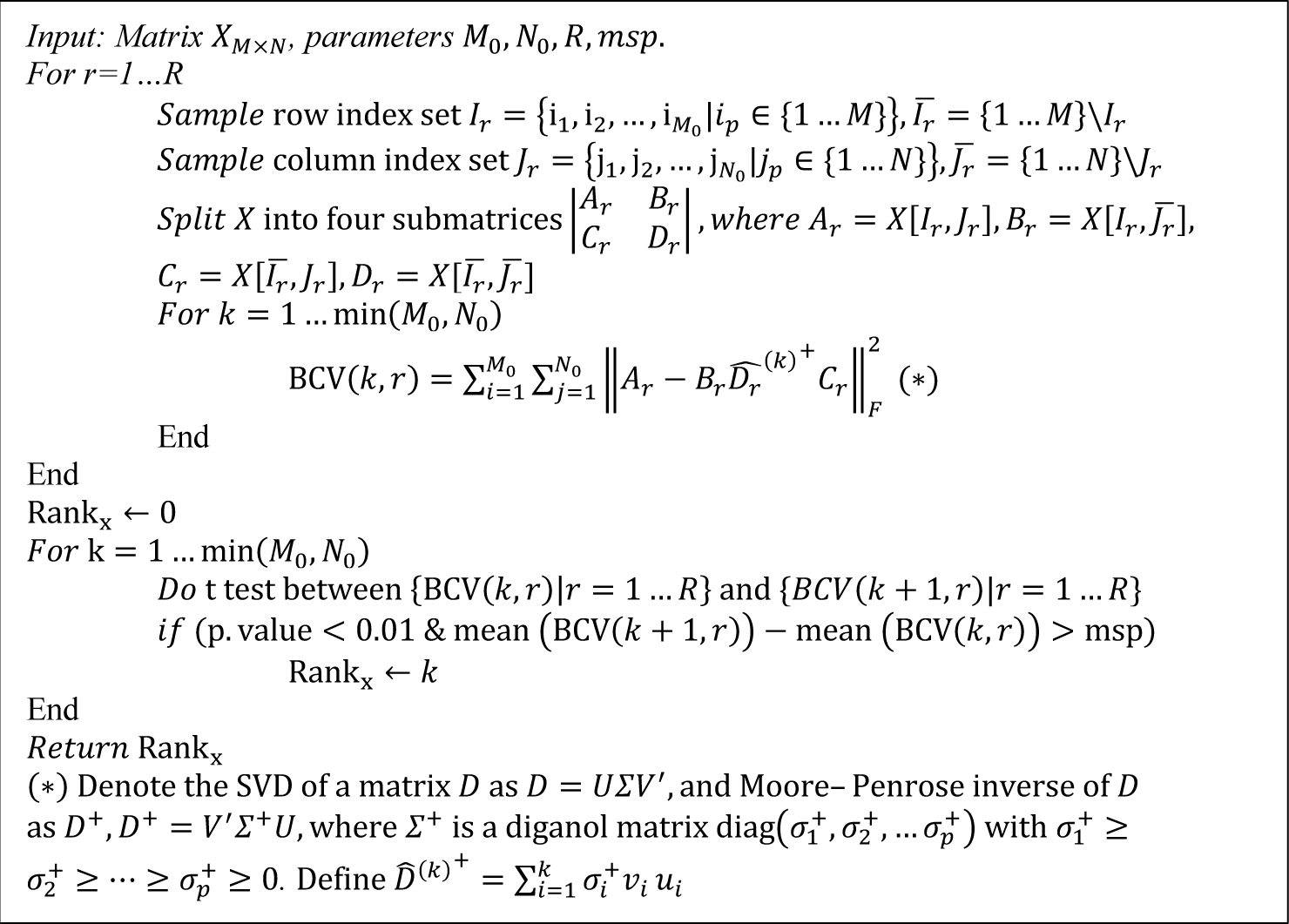

After running the rank-1 module detection on all the training bulk tissue datasets, those genes commonly identified in the rank-1 modules in more than 40% (70%) data sets were selected as core (stringent) markers. The list of stringent marker sets was derived with more stringent criterion, which is particularly useful for the analysis of small sample sized target data. Core markers of cells in central nervous systems were identified by a similar approach on the brain training tissue datasets. Due to the limitation of hematopoietic system tissue training data, its core markers were selected as the genes specifically over expressed in each hematopoietic cell type, by using the criteria: the gene’s expression level is above 10% quantile in one cell type and below 50% in the other cell types. Complete lists of selected core and stringent marker sets were given in **Supplementary Table S1**.

### Estimation of cell proportion

Two methods were utilized to estimate cell proportion: (1) SVD based computation. With cell type specific markers derived, the first row base of the gene expression profile of the marker genes is directly utilized as an estimation of the cell proportion, which can be directly computed by SVD. (2) Constraint NMF based computation. With the number of identifiable cell types and cell type specific markers identified, the signature matrix 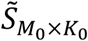 and proportion matrix 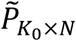can be estimated by minimizing the following objective function:

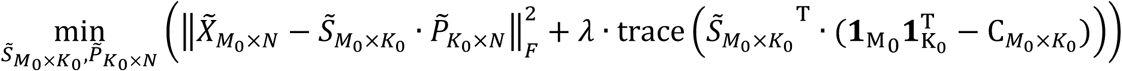

where 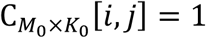 if gene *i* is marker of the cell type *j*, and 0 otherwise. *λ* is the hyper parameter. In this study, we tuned *λ* by using single cell data simulated tissue data. *λ*=10 is empirically utilized in the analysis.

### Explanation score and Comparison with state-of-the-arts methods

An explanation score (ES) was utilized to evaluate the goodness that each marker gene’s expression is fitted by the predicted cell proportions:

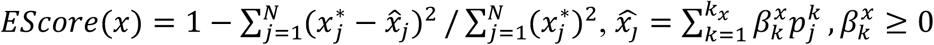

Where 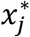 is the observed expression of marker gene *x* in sample *j*, 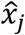 is the explainable expression by cell proportions, obtained by a non-negative regression *x* on the predicted proportion 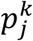,*k* = 1 … *k*_*x*_. Here, *k*_*x*_ represents the number of cell types that express *x*, and 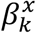 are the non-negative regression parameters. Intuitively, with correctly selected marker genes, the marker gene’s expression can be well explained by the predicted proportions of the cell types that express the gene. Hence, a high ES score is a necessary but not sufficient condition for correctly selected marker genes and predicted cell proportion.

### Data used in this study

#### Bulk cell training data sets

for mouse blood, solid cancer and inflammatory tissue microenvironment, we retrieved 116 datasets of sorted mouse cells of 12 selected cell types, totaling 1106 samples from GEO database. For mouse brain tissue microenvironment, we collected 2130 bulk cell samples of the nine selected cell types in central nerve systems. For mouse hematopoietic microenvironment, two datasets were available that cover 14 hematopoietic cell types. All the bulk cell training data were generated by the Affymetrix GeneChip Mouse Genome 430 2.0 Array platform and normalized with MAS5 method [32]. Samples of the same cell type were further merged together with batch effect removed using Combat [33].

#### Single Cell RNA-sequencing data

One mouse melanoma scRNAseq data set (6638, 9) was acquired from the Human Cell Atlas database [34]. Three scRNA-seq datasets of lung (4485, 12), pancreas (4405, 8), and small intestine (4764, 10) and two sets of brain tissue (3679, 7 and 1099, 6) were accessed from Mouse Cell Atlas (MCA) data portal [35]. The two numbers in the parenthesis indicate the number of cell samples and cell types of each data set. We specifically selected the cells with UMI more than 500 to exclude low quality cells. Cell labels were either provided in the original data or curated using Seurat v3 with cell type specific genes [36, 37].

#### Training tissue data from cancer and blood

33 cancer tissue datasets of 9 cancer types generated by four popular experimental platforms were collected, namely Illumina HiSeq 2000 Mus musculus, Affymetrix Mouse Genome 430 2.0 Array, Illumina HiSeq 2500 Mus musculus and Affymetrix Mouse Genome 430A 2.0 Array from GEO database. Each data set has at least 15 samples. We didn’t consider datasets from immunodeficient mouse, mouse cell lines, and PDX models, as only real cancer or blood micro-environment is considered. A data set of liver tissue collected from 31 mouse strains (GSE55489) were utilized to evaluate the variation of cell type specific markers through different mouse strains [26].

#### Brain tissue data

14 datasets of mouse brain tissues generated by two experimental platforms, namely Illumina HiSeq 2500 Mus musculus and Affymetrix Mouse Genome 430 2.0 Array were collected from Gene Expression Omnibus. Datasets were split into sub data sets of different brain regions. Each data set has at least 40 samples. The complete training data information are available in **Supplementary Table S3**.

#### Hematopoietic System tissue and FACS data

We generated a RNA-seq data set with matched FACS data of bone marrow cells isolated from the hind limbs of C57BL/6, Tet2-/-Flt3ITD, DNMT3A-/-Flt3ITD, and DNMT3A-/-Tet2-/-Flt3ITD mice (n=3 for each group). RNA (600 ng/ sample) was used to prepare single indexed strand specific cDNA library using TruSeq stranded mRNA library prep kit (Illumina). The library prep was assessed for quantity and size distribution using Qubit and Agilent 2100 Bioanalyzer. The pooled libraries were sequenced with 75bp single-end configuration on NextSeq500 (Illumina) using NextSeq 500/550 high output kit. The quality of sequencing was confirmed using a Phred quality score. The sequencing data was next assessed using FastQC (Babraham Bioinfomatics, Cambridge, UK) and then mapped to the mouse genome (UCSC mm10) using STAR RNA-seq aligner [38], and uniquely mapped sequencing reads were assigned by featureCounts. The data were normalized to RPKM. FACS data were collected from same biological prep by IU School of Medicine Flowcytometry Core. Hematopoietic stem cells were identified by lineage negative, C-Kit high and Sca1 high cells, general myeloid progenitor cells were identified by Cd34 and Cd16/32 high cells, mature myeloid cells were identified by Gr1 and Cd11b high cells, and PreB cells were identified by B220 and SSC-A high cells.

### Generation of simulated bulk tissue data from scRNA-seq data

Cell types in each scRNA-seq data were labeled by the cell clusters provided in the original works or by using Seurat pipeline with default parameters. Detailed information of the scRNA-seq data and cell type annotation is given in **Supplementary Table S3**. For each data set, we simulate bulk tissue data by: (1) removing insignificantly expressed genes, (2) randomly generate the proportion of each cell type, called true proportion in this paper, that follows a Dirichlet distribution, and (3) draw cells randomly from the cell pool with replacement according to the cell type proportion, and sum up the expression values of all cells to produce a pseudo bulk tissue data. The insignificant expressed genes were identified by left truncated mixture Gaussian model [39, 40]. The Dirichlet distribution matrix was generated with R package “DirichletReg” [41].

## SUPPLEMENTARY DATA

Supplemental Information can be found in supplementary files

## ACKNOWLEDGEMENTS

C.Z thank Mr. Siyuan Qi from Indiana University School of Medicine for his help in the early stage of this work. C.Z and S.C thank the Indiana University Center for Medical Genomics for their support of this project.

## FUNDING SUPPORTS

This work was supported by the National Science Foundation Div Of Information & Intelligent Systems (No. 1850360). This work was also supported by an R01 award #1R01GM131399-01 from the National Institute of General Medical Sciences. This work was also supported by the Showalter Young Investigator Award from Indiana CTSI.

## CONFLICT OF INTEREST

There are no conflicts to declare.

## Notes

### Competing Interest Statement

The authors have declared no competing interest.

