## Supplementary Notes for "SSMD: A semi-supervised approach for a robust cell type identification and deconvolution of mouse transcriptomics data"

[1]Department of Medical and Molecular Genetics and Center for Computational Biology and Bioinformatics, [4]Department of Pediatrics, [5]Department of Biostatistics, Indiana University School of Medicine, Indianapolis, IN,46202, USA. [2]Department of BioHealth Informatics, Indiana University-Purdue University Indianapolis, Indianapolis, IN, 46202, USA. [3]Department of Electrical and Computer Engineering, Purdue University, Indianapolis, IN, 46202, USA

### SUPPLEMENTARY NOTES

**Benchmark of SSMD with MUSIC on different synthetic data sets.** We selected two types of method to compare with, namely (1) the ones with fixed gene signatures such as ImmunCC, CIBERSORT and TIMER, and (2) the ones enabling training of context specific signature from scRNA-seq data, such as MUSIC. We have demonstrated SSMD beat the first type of methods in the Results (Fig 2). We considered two-fold of comparisons, i.e. compare SSMD with MUSIC (i) on the synthetic tissue data simulated by using scRNA-seq data and the same scRNA-seq as the training data for MUSIC, and (2) synthetic tissue data with using a different scRNA-seq data set as the training data for MUSIC, where (1) is an ideal condition for MUSIC and (2) tests the robustness of MUSIC. As expected, under the condition of (1) MUSIC achieved very good performance while the prediction accuracy of SSMD is only slightly below MUSIC (**Fig 1 of Supplementary Notes**). Under the condition of (2), we conducted the experiment by using different scRNA-seq data for tissue data simulation and marker training in MUSIC, as detailed in the **Table1 of Supplementary Notes**. We identified the averaged prediction accuracy of MUSIC is averaged cor=0.58, which is significantly lower than the averaged cor=0.94 for SSMD (**Fig 2 of Supplementary Notes**) under the condition (2). In addition, we have observed a large disparity of the cell type specific markers among different strains in a liver tissue data set. To the best our knowledge, the availability of scRNA-seq data is far less than covering most tissue and disease types in major mouse strains. Hence, we consider SSMD, which requires less training data, is more ideal for a broader application.

**Table 1 of Supplementary Notes**

| | SSMD | Tissue type |
|---|---|---|
| Melanoma | 0.9146 | Inflammatory |
| Lung | 0.9439 | Inflammatory |
| Pancreas | 0.9501 | Inflammatory |
| Small Intestine | 0.9121 | Inflammatory |
| Brain1 | 0.9730 | Central Nervous System |
| Brain2 | 0.9459 | Central Nervous System |
| **Mean** | **0.9399** | |

| | MUSIC | MUSIC Training |
|---|---|---|
| Melanoma | 0.9966 | Melanoma |
| Lung | 0.9924 | Lung |
| Pancreas | 0.9953 | Pancreas |
| Small Intestine | 0.9712 | Small Intestine |
| Brain1 | 0.9970 | Brain1 |
| Brain2 | 0.9961 | Brain2 |
| **Mean** | **0.9914** | |

| | Noisy MUSIC | MUSIC Training |
|---|---|---|
| Melanoma | 0.6155 | Lung |
| Lung | 0.5999 | Small Intestine |
| Pancreas | 0.8011 | Lung |
| Small Intestine | 0.5548 | Pancreas |
| Brain1 | 0.4549 | Brain2 |
| Brain2 | 0.4748 | Brain1 |
| **Mean** | **0.5835** | |

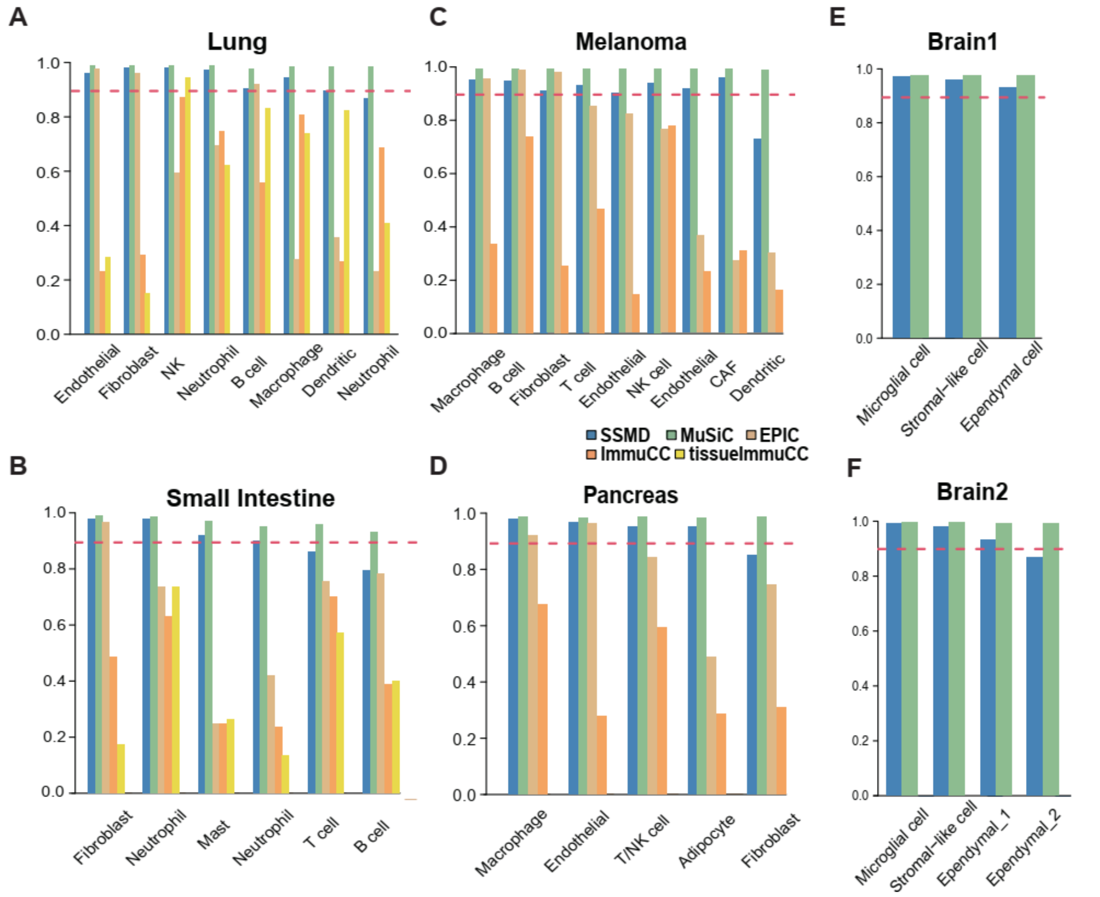


**Fig 1 of Supplementary Notes.** Method evaluation on scRNA-seq simulated tissue data. (A-D) Correlation between true and predicted cell proportions in the simulated Lung (A), Pancreas (B), Small Intestine (C), and Mouse Melanoma (D) tissue data. The x-axis represents cell type and y axis represents prediction accuracy. Predictions made by SSMD, MUSIC, EPIC, ImmuCC and tissue-ImmuCC were dark blue, green, light orange, orange and yellow colored, respectively. The red dash line represents the 0.9 correlation cutoff. (E-F) Correlation between true and predicted cell proportions in the two simulated brain tissue data. The MUSIC prediction was made by using the same scRNA-seq data for both tissue data simulation and marker training.

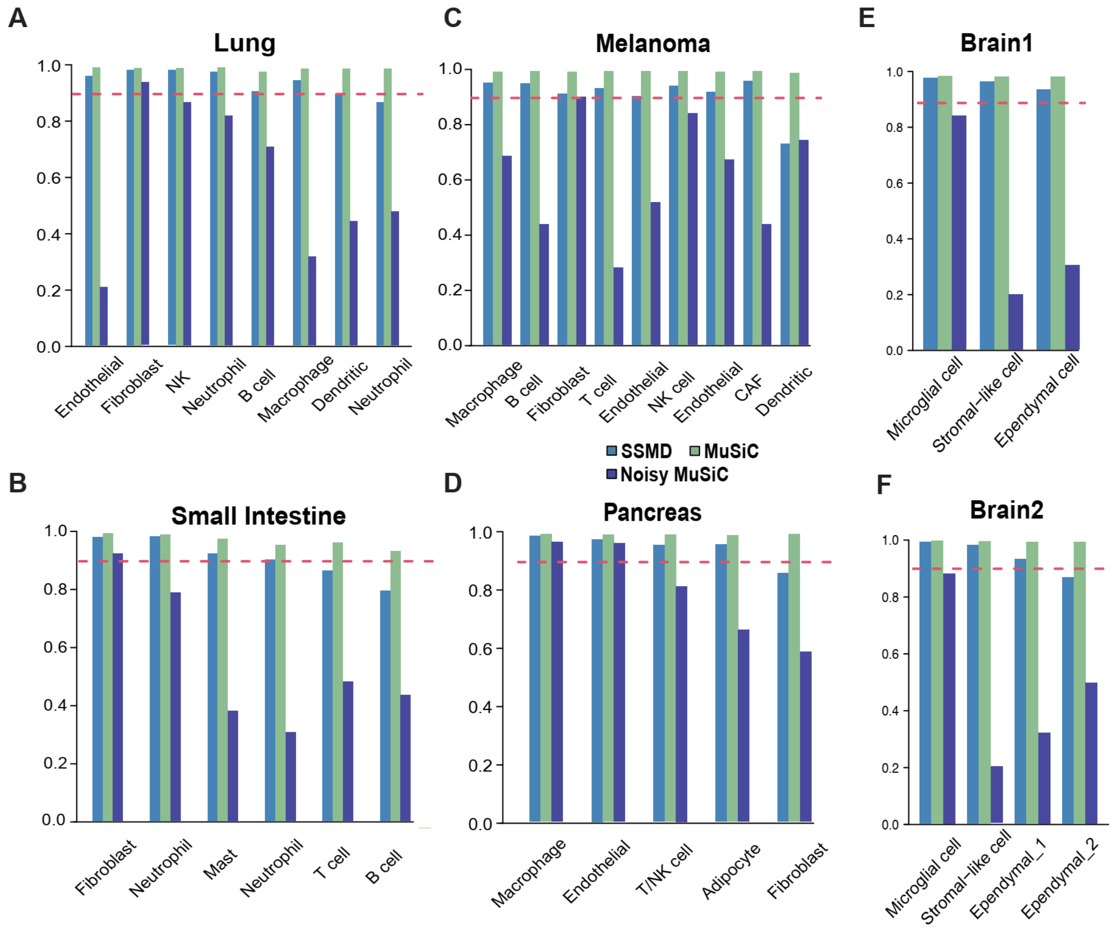


**Fig 2 of Supplementary Notes.** Method evaluation on scRNA-seq simulated tissue data. (A-D) Correlation between true and predicted cell proportions in the simulated Lung (A), Pancreas (B), Small Intestine (C), Mouse Melanoma (D), and Brain (E-F) tissue data. The x-axis represents cell type and y axis represents prediction accuracy. Predictions made by SSMD, MUSIC (condition 1), and noisy MUSIC (condition 2) were blue, green, and dark blue colored, respectively. The MUSIC prediction was made by using the same scRNA-seq data for both tissue data simulation and marker training. The noisy MUSIC prediction was made by using different scRNA-seq data for tissue data simulation and marker training, as detailed in the Table 1 of Supplementary Notes.

### SUPPLEMENTARY FIGURES AND TABLES

#### Supplementary Tables

Supplementary Table S1. Cell type specific core marker list trained by SSMD.

Supplementary Table S2_1. Proportions counted by FACS and predicted by SSMD of the hematopoietic tissue data.

Supplementary Table S2_2. Cell type specific markers of the hematopoietic tissue data

Supplementary Table S3. Training tissue data information.

Supplementary Table S4. Proportion of different cell types in simulated bulk tissue data. And correlations between predicted proportion made by different method VS the ground truth.

#### Supplementary Figures

**Supplementary Figure S1.** Detailed analysis pipeline of SSMD. (A) Analysis pipeline of the core marker training procedure. (B) Analysis pipeline of the deconvolution procedure.

**Supplementary Figure S2.** SSMD web server and utilization guideline. (A) A screen shot of SSMD webserver. (B) A best practice guide for SSMD webserver and package.

**APPENDIX**

**Theoretical bases of "transcriptomically identifiable" cell types:**

Below we list the necessary theoretical derivations of the mathematical condition for "transcriptomically identifiable" cell types. Complete derivations are available at https://github.com/xiaoyulu95/SSMD. The goal of the theoretical bases listed below is to support that the "rank-1" pattern of gene markers is a necessary but non-sufficient condition for transcriptomically identifiable cell types. If this condition does not hold, there is no computational method cannot guarantee the correlation between the predicted and true proportion of the cell type is asymptomatically converge to 1 or sufficiently high, i.e. the cell type is not computational identifiable.

### *1. Notations:*

A tissue expression profile could be thought as generated by pooling mRNAs from all of its cells, the total mRNA abundance in one tissue sample can then be considered as the following model:

$$X^i_{M\times 1} = \sum_{c \in Tissue\ i} x^{c,i}_{M\times 1} + Z^i_{M\times 1} \ (1)$$

where $X^i_{M\times 1}$ denotes the mRNA abundance of $M$ genes in the $ith$ tissue sample and $x^{c,i}_{M\times 1}, c \in Tissue\ i$ represents the mRNA profile of the $c$th cell in the tissue, $Z^i_{M\times 1}$ represents the extracellular mRNA abundance. Assume there are $C^i$ cells in tissue $i$, in which $C^i_k$ number of cell are of type $k, k = 1 \dots K,$ in tissue $i$, such that $\sum_{k=1\dots K} C^i_k = C^i$. Then, equation (1) can be written as:

$$X^i_{M\times 1} = \sum_{k=1}^{K} \sum_{j=1}^{C^i_k} x^{k,i}_{M\times 1,j} + Z^i_{M\times 1} \ (2)$$

where $x^{k,i}_{M\times 1,j}$ denotes the gene expression profile of the $j$th cell of type $k$ in the tissue sample $i$. The exact population and proportion of cell type $k$ in the sample $i$ are $C^i_k$ and $C^i_k/C^i$, respectively. Denote $P^i_k = C^i_k/C^i$.

Caused by the PCR amplification and limited by the sequencing amount, denoted $\tilde{X}^i_{M\times 1}$ as the measured gene expression profile from $X^i_{M\times 1}$, which is first normalized by the observed gene expression signal through different tissue samples. Denote $|X_{M\times 1}|_{L_1}$ as the sum of all elements in $X_{M\times 1}$, i.e., the $L_1$ norm, and we have

$$\tilde{X}^i_{M\times 1} = \mathrm{B_i} \cdot \frac{\sum_{k=1}^{K} \sum_{j=1}^{C^i_k} x^{k,i}_{M\times 1,j} + Z^i_{M\times 1} + \epsilon_{0,i}}{\sum_{k=1}^{K} \sum_{j=1}^{C^i_k} \left| x^{k,i}_{M\times 1,j} \right|_{L1} + |Z^i_{M\times 1}|_{L1}} + \epsilon_{1,i} \quad (3)$$

, where $\mathrm{B_i}$ denotes the total observed gene expression signal in sample i, an analog of the total sequenced counts of RNA-seq data, $\epsilon_{0,i}$ and $\epsilon_{1,i}$ represent the library preparation (or sample preparation) error and measurement error respectively.

Let $\bar{x}^{k,i}_{M\times 1}$ denote the mean expression profile of the cells of cell type $k$ in the $i$th tissue sample, we have

$$\tilde{X}^{i}_{M\times 1} = B_{\mathrm{i}} \cdot \frac{\sum_{k=1}^{K}\sum_{j=1}^{C_k^i} x^{k,i}_{M\times 1,j} + Z^{i}_{M\times 1} + \epsilon_0}{\sum_{k=1}^{K} C_k^i \left|\bar{x}^{k,i}_{M\times 1}\right|_{L_1} + |Z^{i}_{M\times 1}|_{L1}} + \epsilon_1 \quad (4)$$

Denote $\bar{B}_i = \frac{\sum_{k=1}^{K} C_k^i \bar{x}^{k,i}_{M\times 1} + |Z^{i}_{M\times 1}|_{L1}}{C^i} = \sum_{k=1}^{K} P_k^i \bar{x}^{k,i}_{M\times 1} + \frac{|Z^{i}_{M\times 1}|}{C^i}$, we further have

$$\tilde{X}^{i}_{M\times 1} = \frac{B_{\mathrm{i}}}{\bar{B}_i}\sum_{k=1}^{K}\frac{\sum_{j=1}^{C_k^i} x^{k,i}_{M\times 1,j}}{C^i} + \frac{B_{\mathrm{i}}}{\bar{B}_i}\frac{Z^{i}_{M\times 1}}{C^i} + \frac{B_{\mathrm{i}}}{\bar{B}_i}\epsilon_0 + \epsilon_1 \quad (5)$$

Equation (5) characterizes the relationships among the gene expression signal from each single cell, sample wise variations in cell type proportion, total observed gene expression signal, extracellular mRNA and errors on the observed tissue level gene expression. Based on (5), we will investigate the impact of different assumptions on estimating $P_k^i$, for the deconvolution of $N$ tissue samples $\tilde{X}_{M\times N} = \{\tilde{X}^{i}_{M\times 1} | i = 1 \dots N\}$.

Historically, deconvolution assumes $\tilde{X}_{M\times N}$ as the following non-negative product form:

$$\tilde{X}_{M_0\times N} = \tilde{S}_{M_0\times K_0} \cdot \tilde{P}_{K\times N} + E, \tilde{S}_{M_0\times K_0} \geq 0, \tilde{P}_{K_0\times N} \geq 0 \quad (6)$$

,where $\tilde{X}_{M_0\times N}$ represents the observed gene expression matrix of $M_0$ selected genes in $N$ tissue samples, and $\tilde{S}_{M_0\times K_0}$, usually unknown, denotes the gene expression signature matrix, and $\tilde{P}_{K_0\times N}$ denotes the proportion matrices of $K_0$ cell types, to be estimated. To estimate $\tilde{P}_{K_0\times N}$, the loss function of a deconvolution problem is usually written as

$$\mathbf{min}\left(\left\|\tilde{\boldsymbol{X}}_{\boldsymbol{M_0\times N}} - \tilde{\boldsymbol{S}}_{\boldsymbol{M_0\times K_0}} \cdot \tilde{\boldsymbol{P}}_{\boldsymbol{K\times N}}\right\|_{\boldsymbol{L2}} + \boldsymbol{penalty} \mid \boldsymbol{constraint}\right) \ (\mathbf{7})$$

, where $\|X\|_{L_2}$ denotes $L_2$ norm of $X$, i.e., sum of square of all elements, and the $penalty$ and $constraint$ vary by methods. Let $\tilde{P}_{K_0\times N}$ be a solution to the optimization problem in (7), then, $E\left(\tilde{X}^{i}_{M_0\times 1} - \tilde{S}_{M_0\times K_0} \cdot \tilde{P}^{i}_{K_0\times 1}\right) \approx 0$. Successful deconvolution using formulation (7) needs to ensure $\tilde{P}_{K_0\times N}$ to be close to the true proportion matrix, $P_{K_0\times N}$, meaning either their absolute deviance is small, or loosely, their Pearson correlation is high.

### 2. *Assumptions*

***Condition I:*** Signature genes in $M_0$ should be properly selected such that enable $E\left(\frac{B_i}{\bar{B}_i}\frac{Z^i_{M_0\times 1}}{C^i}\right)=0$, $\sum_{k=K_1+1}^{K_2} E\left(\frac{B_i}{\bar{B}_i}x^{k,i}_{M_0\times 1,j}P^i_k\right)=0$, and additionally, genes of which $\tilde{S}^t_{M_0\times 1}\neq 0, t\in\{K_2+1,\ldots,K_3\}$ should be none or lowly expressed in the tissue data.

***Condition II.1 (non-negative regression formulation).*** For $K_1$ cell types that both present in the tissue samples and are considered in the $\tilde{S}_{M_0\times K_1}$ matrix, if $\hat{S}^i_{M_0\times 1}\perp \hat{S}^j_{M_0\times 1}|i,j\in\{1\ldots K_1\}, i\neq j$, and if for any $\hat{S}^i_{M_0\times 1}$, the best non-negative regression of $\hat{S}^i_{M_0\times 1}=\sum_{j=\{1\ldots K_1\}\backslash\{i\}}\beta_j\hat{S}^j_{M_0\times 1}$ is $\beta_j=0, j=\{1\ldots K_1\}\backslash\{i\}$, i.e. $\hat{S}^i_{M_0\times 1}$ is not positively associate with any non-negative linear combination of $\hat{S}^j_{M_0\times 1}, j=\{1\ldots K_1\}\backslash\{i\}$, and $cor\left(\hat{S}^k_{M_0\times 1},\tilde{S}^k_{M_0\times 1}\right)>0, k=1\ldots K_1$ and $cor\left(\hat{S}^i_{M_0\times 1},\tilde{S}^j_{M_0\times 1}\right)<0, i\neq j$, then $cor\left(\tilde{P}^{i=1\ldots N}_k, P^{i=1\ldots N}_k\right)=1, k=1\ldots K_1$.

***Condition II.2 (general regression formulation).*** For the $K_1$ cell types that both present in the tissue samples and are considered in the $\tilde{S}_{M_0\times K_1}$ matrix, if $\hat{S}^i_{M_0\times 1}\perp \hat{S}^j_{M_0\times 1}|i,j\in\{1\ldots K_1\}, i\neq j$, and for any $i$, $cor\left(\hat{S}^i_{M_0\times 1},\tilde{S}^i_{M_0\times 1}\right)>0, \forall i$ and $\hat{S}^i_{M_0\times 1}\perp \hat{S}^j_{M_0\times 1}, \forall\, i\neq j$, then $cor\left(\tilde{P}^{i=1\ldots N}_k, P^{i=1\ldots N}_k\right)=1, k=1\ldots K_1$.

***Condition II.3 (necessary but not sufficient condition for un-orthogonal cell type signature expression).*** Suppose $\exists\, i,j\in\{1\ldots K_1\}$, s.t. $\hat{S}^i_{M_0\times 1}$ is not orthogonal to $\hat{S}^j_{M_0\times 1}$, then the necessary but not sufficient condition for $cor\left(\tilde{P}^{i=1\ldots N}_k, P^{i=1\ldots N}_k\right)=1, k=1\ldots K_1$ is that $\tilde{P}_{K_1\times N}$ provides a good non-negative bases for $\tilde{X}_{M_0\times N}$, meaning $\tilde{P}_{K_1\times N}$ and its corresponding $\tilde{S}_{M_0\times K_1}$ is (close to) an optimal solution to NMF of $\tilde{X}_{M_0\times N}$.

It is noteworthy that if the ***Condition II.1*** or ***Condition II.2*** holds, and the matrix rank of $\tilde{X}_{M_0\times N}$ is $K_1$, **the predicted** $\tilde{P}^{i=1\ldots N}_k$ is the unique solution for the NMF of $\tilde{X}_{M_0\times N}$.

***Condition III.1 (Uniqueness of solution).*** For the deconvolution of a tissue data generated by $\tilde{X}_{M_0\times N} = \hat{S}_{M_0\times K_1} \cdot P_{K_1\times N}$, the NMF problem $\left\|\tilde{X}_{M_0\times N} - \bar{S}_{M_0\times K_1} \cdot \bar{P}_{K_1\times N}\right\|$ has a unique solution $\hat{S}_{M_0\times K_1}$ and $P_{K_1\times N}$ only if $\hat{S}^i_{M_0\times 1} \perp \hat{S}^j_{M_0\times 1} | i,j \in \{1 \ldots K_1\}, i \neq j$.

***Condition III.2 (Signature matrix constraint).*** A sufficient condition for the constraint NMF problem $\left\|\hat{S}_{M_0\times K_1} \cdot P_{K_1\times N} - \bar{S}_{M_0\times K_1} \cdot \bar{P}_{K_1\times N}\right\| - \lambda \cdot trace(\bar{S}^T_{M_0\times K_1} \cdot \tilde{S}_{M_0\times K_1})$ has a unique solution $\hat{S}_{M_0\times K_1}$ if $\bar{S}_{M_0\times K_1} = \hat{S}_{M_0\times K_1}$ minimizes $trace(\bar{S}^T_{M_0\times K_1} \cdot \tilde{S}_{M_0\times K_1})$.

***Condition III.2*** is suggesting that among the multiple solutions that minimize $\left\|\tilde{X}_{M_0\times N} - \bar{S}_{M_0\times K_1} \cdot \bar{P}_{K_1\times N}\right\|$, we should select the one that most resembles the prior $\tilde{S}_{M_0\times K_1}$.

#### *3. Theorems*

***Definition I. Transcriptomically identifiable cell types***. For a given transcriptomics data set $X_{M\times N}$ and a deconvolution method, a cell type $k$ defined as "transcriptomically identifiable" if its true proportion $P^k_{1\times N} \triangleq \left\{\frac{C^1_k}{C^1}, \frac{C^2_k}{C^2}, \ldots \frac{C^N_k}{C^N}\right\}$ and estimated proportion $\tilde{P}^k_{1\times N}$ are perfectly linearly correlated, i.e. $cor\left(P^k_{1\times N}, \tilde{P}^k_{1\times N}\right) = 1$

***Definition II (explanation score, ES).*** The explanation score of gene $g$ is defined by

$$ES_g \triangleq min \frac{\sum_{i=1}^{N}\left(\tilde{X}_{M_0\times N}[g,i] - \sum_{k\in CS_g} \beta_k \hat{P}^i_k\right)^2}{\sum_{i=1}^{N}\left(\tilde{X}_{M_0\times N}[g,i] - mean(\tilde{X}_{M_0\times N}[g,])\right)^2} \mid \beta_k \geq 0.$$

***Theorem 1.1*** For a supervised method, the $K_1$ cell types in formula (5) are ***transcriptomically identifiable*** if and only if the ***Condition I*** and ***Condition II.1*** *or* ***Condition II.2*** strictly hold, and the $E(\epsilon_0)$ and $E(\epsilon_1)$ can be ignored.

***Theorem 1.2*** For a unsupervised NMF based method, the $K_1$ cell types in formula (5) are ***transcriptomically identifiable*** if and only if the ***Condition I*** and ***Condition III.1*** strictly hold, and the $E(\epsilon_0)$ and $E(\epsilon_1)$ can be ignored.

***Corollary 1.3*** If the $K_1$ cell types in formula (5) are ***transcriptomically identifiable*** by a supervised method, they are also ***transcriptomically identifiable*** by an unsupervised method.

***Theorem 2.1 (strong identifiability).*** For a given tissue transcriptomics data $X_{M\times N}$, a cell type $k$ is transcriptomically identifiable in all the samples, if and only if (1) there are genes $G_k$ uniquely expressed by cell type $k$, s.t. $E(X_{G_k\times T})$ is a rank one matrix, for $\forall\, T$ is a subset of $\{1, \dots, N\}$.

***Theorem 3 (Necessary Condition)***. If the gene $g$ is a marker gene of several transcriptomically identifiable cell types $CS_g$ in all the samples, $\lim_{N\to\infty} ES_g = 0$.
